# Functional tagging of endogenous proteins and rapid selection of cell pools (Rapid generation of endogenously tagged *piwi* in ovarian somatic sheath cells.)

**DOI:** 10.1101/2020.12.18.423517

**Authors:** Celine Marlin Andrews, Parthena Konstantinidou, Pavol Genzor, Daniel Stoyko, Alexandra R. Elchert, Leif Benner, Sushil Sobti, Esther Y. Katz, Qingcai Meng, Astrid D. Haase

**Affiliations:** National Institutes of Diabetes and Digestive and Kidney Diseases, National Institutes of Health, Bethesda, MD; Department of Biochemistry, School of Medicine, University of Patras, 26504 Patras, Greece; Department of Biology, Johns Hopkins University, Baltimore, MD, 21218, USA

**Author notes:** These authors contributed equally (Celine Marlin Andrews, Parthena Konstantinidou, Pavol Genzor, Daniel Stoyko).

## Abstract

The combination of genome-editing and epitope tagging provides a powerful strategy to study proteins with high affinity and specificity while preserving their physiological expression patterns. However, stably modifying endogenous genes in cells that do not allow for clonal selection has been challenging. Here, we present a simple and fast strategy to generate stable, endogenously tagged alleles in a non-transformed cell culture model. At the example of *piwi* in *Drosophila* ovarian somatic sheath cells, we show that this strategy enables the generation of an N-terminally tagged protein that emulates the expression level and subcellular localization of the wild type protein and forms functional Piwi-piRNA complexes. We present a concise workflow to establish modified cells, characterize the edited allele and probe the function of the tagged protein.

## INTRODUCTION

Epitope tags provide high specificity and high affinity handles for our favorite genes. Optimized tags for visualization, purification and even functional manipulation can be used to track the localization of proteins, identify interaction partners and even induce degradation (1). Excellent tools are readily available and there is no need to engage in the laborious and expensive process of generating and optimizing antibodies for individual proteins (2). Epitope tags would be perfect, if they could be added to any protein without changing its expression and function. N-terminal or C-terminal tagging strategies, flexible linkers and small or globular tags provide a variety of combinations that can be adjusted to generate functionally tagged targets (1). However, maintaining the expression level and regulation of the endogenous genes has been challenging. Before CRISPR-assisted genome editing, adding a tag directly to an endogenous gene has only been practical in embryonic stem cells that support a high degree of homologous recombination, and involved the selection and clonal expansion of a single modified cell (2–4). Modern CRISPR strategies have expanded the repertoire of modifiable cell types but still mostly rely on the identification and clonal expansion of a single modified cell to establish a stable clonal cell line (5–7). Generation of mutant clones is generally the most time-consuming and laborious step in CRIPSR-based genome editing, and some cell culture systems do not at all support efficient clonal expansion.

Such a delicate cell culture system is *Drosophila* ovarian somatic sheath cells (OSC), a unique *ex vivo* model for a specialized small RNA silencing pathway that restricts mobile genetic elements, transposons, to guard genome integrity of germ cells (8) (Fig. 1A). PIWI-interacting small RNAs (piRNAs) and their PIWI protein partners form functional piRNA silencing complexes that recognize transposon transcripts by sequence complementarity and induce silencing at transcriptional and post-transcriptional level (9–11). OSC were established from adult *Drosophila* ovaries initially as part of a co-culture system for germ cells (12). OSC reflect follicle cells of the germ cell niche and operate a Piwi-only piRNA pathway that induces transcriptional silencing of endogenous retroviruses (8). Piwi is one of three PIWI-clade Argonaute proteins in Drosophila (13). Mature Piwi-piRNA complexes transition to the nucleus, target nascent transposon transcripts and recruit the H3K9 histone methyl transferase Eggless/dSetDB1 to establish epigenetic restriction (14). Upon knock-down of *piwi*, restriction of various transposons is lost, and cells die (15, 16). The OSC represent the only *ex vivo* system to study transcriptional silencing by Piwi-piRNA complexes and has become an invaluable tool for piRNA biology. Here, we developed a strategy for epitope-tagging of endogenous proteins, rapid selection of edited cell pools, and established an endogenously FLAG-HA tagged *piwi* allele (eFH-*piwi*) in ovarian somatic sheath cells (OSC) that emulates the expression and function of wild type *piwi*.

## MATERIALS AND METHODS

### Design and preparation of sgRNAs

The sgRNA sequence was designed using the GuideScan algorithm (17). sgRNAs with no off-targets effects were chosen based on their proximity to the start codon. Complementary oligonucleotides, each with appropriate 5’ overhangs (ps12_sgRNA and pas13_sgRNA, see supplementary table), were purchased from integrated DNA technologies (IDT). These oligos were annealed and cloned into pU6-BbsI-chiRNA (addgene #45946) after linearization with BbsI.

### Design and generation of the donor plasmid for homologous repair

The donor plasmid was designed to contain an intronic selection cassette that allows the expression of the puromycin resistance gene from the opposite genomic strand. The splice donor and splice acceptor sites were modelled after the Drosophila MHC gene (exon 17: donor and exon 19: acceptor). A Kozak sequence followed by a start codon (AATCAAA_ATG) were placed after the intron in frame with a combined 3xFLAG-3xHA (FH) tag. The tags were spaced by flexible linkers (GSS). The intron contained a puromycin resistance gene driven by an Actin 5C promoter on the opposite genomic strand and inverted in orientation to avoid interference. The entire donor cassette containing the intron and the FH-tag were flanked by BbsI restriction sites that allow for insertion of homology arms. Homology arms, of about 750 bp, were amplified from OSC genomic DNA using primers 1-4 and cloned into the donor plasmid using HiFi DNA Assembly (NEBuilder). (see supplementary table for primers and plasmids)

### Cell culture and transfection of ovarian somatic sheath cells (OSC)

OSC were cultured according to the initial instructions (12). The donor plasmid, the sgRNA plasmid and a plasmid expressing the Cas9 nuclease were transfected using the Xfect Transfection Reagent (631318, Takara). A Cas9 expression plasmid without a puromycin resistance was generated by removing puromycin from pAc-sgRNA-Cas9 (addgene# 49330). OSC cells seeded in 10 cm dishes, were transfected with a total of 30 μg of the plasmids after reaching 40-50% confluency. The transfection mixture (plasmids, Xfect polymer, Xfect buffer) was added to the cells after replacing the complete medium (12) with medium lacking fly extract. The cells were incubated at 25° C for three hours after which, the minimal medium containing the transfection mixture was replaced with complete medium. To prevent nonhomologous end joining the cells were treated with SCR7 (Selleckchem, S7742) at 5μM/mL upon replacement of complete medium, and again at 24 hours post transfection. Starting at 48 hours post transfection, cells were treated with puromycin at 2μg/mL. During antibiotic selection, non-edited cells died within 3-5 days and stably edited cells, OSC:*eFH-piwi,* replenished the population within 2-3 weeks.

### Verification of genome editing by genomic PCR

Genomic DNA was extracted from *OSC:eFH-piwi* and wild type (wt) OSC using gDNA kit (Zymo Research). PCR amplification was done using Q5 High-Fidelity DNA Polymerase (New England Biolabs, M0491). To amplify the transcript from the modified allele, we combined universal primers that recognize sequences in the Flag-HA tag and gene-specific primers that recognize the genomic *piwi* locus. The gene-specific primers were designed to reside outside the homology arms, primer ps1_piwi and primer pas2_piwi. Primers ps1_piwi and pas2_piwi were also coupled with primer pas3_intron and primer ps4_flag, respectively. PCR products were separated by a 1% Agarose gel electrophoresis and visualized using GelRed.

### Characterization of the mature edited mRNA

Total RNA was extracted from OSC:*eFH-piwi* and wild type (WT) OSC using Trizol and Direct-zol RNA MiniPrep kit (Zymo Research, R2051). Complementary DNA (cDNA) was generated using the SuperScript IV (Thermo Fisher Scientific, 18090010) reverse transcription reagents and oligo dT_20_ primers. For PCR amplification of the transcript from the modified allele, we combined universal primers that recognize sequences in the Flag-HA tag (pas5_flag, pas7_tag) and gene-specific primers that recognize the genomic *piwi* locus (ps6_UHA). PCR products were separated by a 1% Agarose gel electrophoresis and visualized using GelRed.

### Protein quantification by western blotting

*OSC:eFH-piwi* were lysed in 100 μl lysis buffer (20 mM Tris-HCl pH 7.4, 250 mM NaCl, 2 mM MgCl_2_, 1% NP-40) supplemented with 1x Protease Inhibitor (Thermo Fisher Scientific, 1861281) and 0.1 U/μL Universal Nuclease (Thermo Fisher Scientific, 88701) and were incubated on ice for 15 minutes. The Cell lysates were centrifuged at full speed for 10 minutes, and the cleared lysate was transferred to new tube. After addition of reducing LDS Sample Buffer (Thermo Fisher Scientific, 84788), the lysates were further denatured at 95°C for 3 minutes, allowed to cool at room temperature and spun. Protein contents were separated through a NuPAGE 4-12% Bis-This Gel (Invitrogen, NP0321BOX). The separated proteins were transferred to a PVDF membrane. The membrane was incubated in Odyssey Blocking Buffer (PBS) (LI-Cor, 927-40100) supplemented with 0.1% Tween for 30 minutes at room temperature. Following blocking, the membrane was incubated with 1:2500 Rabbit anti-Piwi polyclonal antibody (18) in blocking buffer at 4°C overnight. The excess of antibody was washed off with TBST three times for 5 minutes and the membrane was incubated with 1:10000 IRdye800 goat anti-Rabbit 2ry antibody (cat No: 92632211, Li-COR) for fifty minutes at room temperature in a dark container. Finally, the membrane was washed with TBST five times for 5 minutes and fluorescence was detected in the Odyssey Infrared Imaging System.

### Determination of subcellular localization by immunofluorescence and microscopy

*OSC:eFH-piwi* were allowed to adhere to concanavalin A treated glass slides overnight. Cells were fixed using 4% PFA, permeabilized with 0.1% Triton x100, and blocked with 3% filtered BSA solution. Samples were washed with PBS following each step. The sample was probed using an Anti-HA High Affinity antibody (Sigma Aldrich, 11867423001) followed by Goat anti-rat IgG (H+L) Alexa Fluor 568 secondary antibody (Thermo Fisher Scientific, A-11077). DAPI was applied at a concentration of 1μg/ml. Slides were sealed with a cover slip using ProLong™ Glass Antifade Mountant (Thermo Fisher Scientific, P36980). Images were taken 24 hours later using an LSM 700 confocal microscope at 100x magnification.

### Purification of Piwi-piRNA and FH-Piwi-piRNA complexes and preparation of associated piRNAs for high-throughput sequencing

PiRNA samples were prepared for wild-type (WT) and the endogenously Flag-HA-tagged Piwi (eFH-Piwi) protein (3 biological replicates each) using WT OSC and *OSC:eFH-piwi* respectively.

OSC cell pellets were dissolved in cold IP buffer (20mM Tris HCl pH7.4, 250mM NaCl, 2mM MgCl_2_, 1% NP40) supplemented with 1x Halt Protease & Phosphatase Inhibitor Cocktail (Thermo Fisher Scientific, 1861281) and were incubated on ice for 10 minutes. Lysates were then centrifuged at 13000g for 15 minutes and the supernatant (input) was used for immunoprecipitation. Lysates of OSC:*eFH-piwi* were incubated with 30μl anti-FLAG M2 magnetic beads (Sigma-Aldrich, M8823). Lysates of WT OSC were incubated with 1μg of Rabbit anti-Piwi polyclonal antibody (18) and 30 μl Surebeads Protein A Magnetic beads (Bio-Rad, 1614013). Immunoprecipitation was performed at 4°C overnight. Next day, the beads were washed three times with IP buffer, one time with high salt buffer (20mM Tris HCl pH7.4, 500mM NaCl, 2mM MgCl_2_, 1% NP40), followed by one wash with IP buffer to remove the excess salt. Immunoprecipitated RNA was recovered with Trizol using the Direct-zol RNA MiniPrep kit (Zymo Research, R2051).

The purified piRNAs were prepared for Illumina sequencing using the general protocol described by Hafner M. et al. (19). In brief, RNAs were ligated to a 29nt long ^32^P-labeled 3’ index adapter and ligation products were recovered from a 12% Urea PAGE gel by extracting 48-58nt long fragments (corresponding to 19-2nt long input small RNAs). Next, 3’-ligated RNA was ligated to the 34nt long 5’ DNA-RNA hybrid adaptor and ligation products (82-92nt long fragments) were recovered from a 10% Urea PAGE gel. piRNA libraries were sequenced using the Illumina HiSeq 3000 and obtaining 50nt single end reads. 5’ and 3’adapters included a total of 10 unique molecular identifiers (UMIs) that allowed for elimination of PCR duplicates during bioinformatic analysis. See supplemental table for adaptor sequences.

### Initial computational processing of raw sequencing data

The raw files (*fastq*) were processed by removing the constant adapter regions (5’ & 3’; *cutadapt v2.3*) and retaining only reads > 19-nucleotides (nt) in length. To remove PCR duplicates, reads were collapsed to unique sequences before removal of UMIs and only reads >=20 nt in length were retained for further analysis. To optimize the file size for downstream analysis, piRNA reads were collapsed by sequence and stored in *fasta* format that retained multiplicity information. Each sequence has *a fasta* header according to the following formula: **SAMPLE_NAME-S[*id*#]M[*abundance#*]**, where *id#* is the unique order of each sequence, and *abundance#* is the number of times each sequence was present. Our raw and processed files (* *UNIQSEQS.fasta*) files are available online (GEO: GSE156058).

### Filters and genome mapping

To remove potential contaminating RNA fragments from high-abundant cellular RNAs, the samples were mapped to ‘structural RNAs’ (tRNA, rRNA, snRNA, snoRNA; UCSC genome browse; dm6 assembly) using STAR aligner (v2.5.2b). The unmapped reads were then mapped to the dm6 genome allowing for up to 100 multimapping positions (STAR, v2.5.2b) (20).

### Data analyses and plotting

Mapped data were analyzed in *R* as follows. The primary alignments (flag = 0 & 16) of perfectly mapping sequences (tag NM = 0) ranging from 18 to 32-nucleotides (nt) in length were extracted from bam files. The abundance information (*M*[*abundance#*]) was retrieved from sequence names. Read length distribution was calculated by first multiplying each sequence by its abundance, then counting the number of reads per size range, and finally dividing counts by total library size. Metagene analyses were performed by considering only uniquely mapping sequences (NH = 1). The sequences were centered at either 5’end (position 1 = 1st nucleotide of piRNA). The region surrounding the center was expanded by 50nt in both directions producing a 100-nt interval. Genomic sequence for each interval was retrieved, duplicated by its abundance, and used to calculate the nucleotide frequency matrix. Annotations by genomic origin were performed only for uniquely mapping sequences (NH = 1). Each sequence was duplicated by its abundance. The genomic positions were determined by intersecting reads first with sense and then with antisense of repeatmasker (dm6 genome; rmsk_te) (21), exon and intron (UCSC; dm6 genome; refgene) annotation files in order. The unannotated sequences were labeled as other. All data were plotted using ggplot package.

Illumina sequencing data are available at GEO: GSE156058. All reagents are available upon request and will be deposited to a publicly available repository after publication.

## RESULTS

### Generation of an endogenously tagged *piwi* allele and rapid selection of stable cell pools

In order to insert an epitope tag into the open reading frame of *piwi*, we designed an sgRNA to target the *piwi* gene close to the translation start codon (ATG). We would like to note that genetic polymorphisms between the OSC and the *Drosophila* reference genome (dm6) could hamper the targeting potential of an sgRNA that is designed based on the reference. Thus, we first tested the genomic target region by sanger sequencing and selected an sgRNA that can efficiently target the OSC genome. Next, we generated a donor construct for homologous repair with the aim to insert a FLAG-HA(FH)-tag in frame with *piwi*’s ATG and to independently expresses a puromycin resistance gene allowing for rapid selection of edited cells (Fig. 1A). To accommodate the puromycin resistance without disrupting *piwi*, we inserted an intron immediate upstream of the FH-tag. The intron contained the puromycin resistance gene driven by a constitutive promoter. Our design aimed to express two independent transcripts from different genomic strands: The first transcript is driven by the endogenous *piwi* promoter and generates a mature mRNA that only differs from the wild type (WT) transcript by an additional exon-exon junction and a FH-tag fused to *piwi*’s open reading frame (ORF). The second transcript is produced from the opposite genomic strand and produces an independent mRNA encoding the puromycin resistance (Fig. 1A). We co-transfected the plasmid expressing the sgRNA and the donor construct together with a Cas9 CRISPR nuclease into OSC and treated the cells with the DNA-Ligase IV inhibitor SCR7 to increase the probability for homologous repair (HR) (22) (Fig. 1B). We started selection of edited cells using Puromycin 48 hours after transfection. Wild type OSC were sensitive to 2μg/ml Puromycin and died within three to five days. Successfully edited cells were resistant to the puromycin treatment and reconstituted a healthy cell population (OSC: *eFH-piwi)* within two to three weeks.

**Figure 1.**
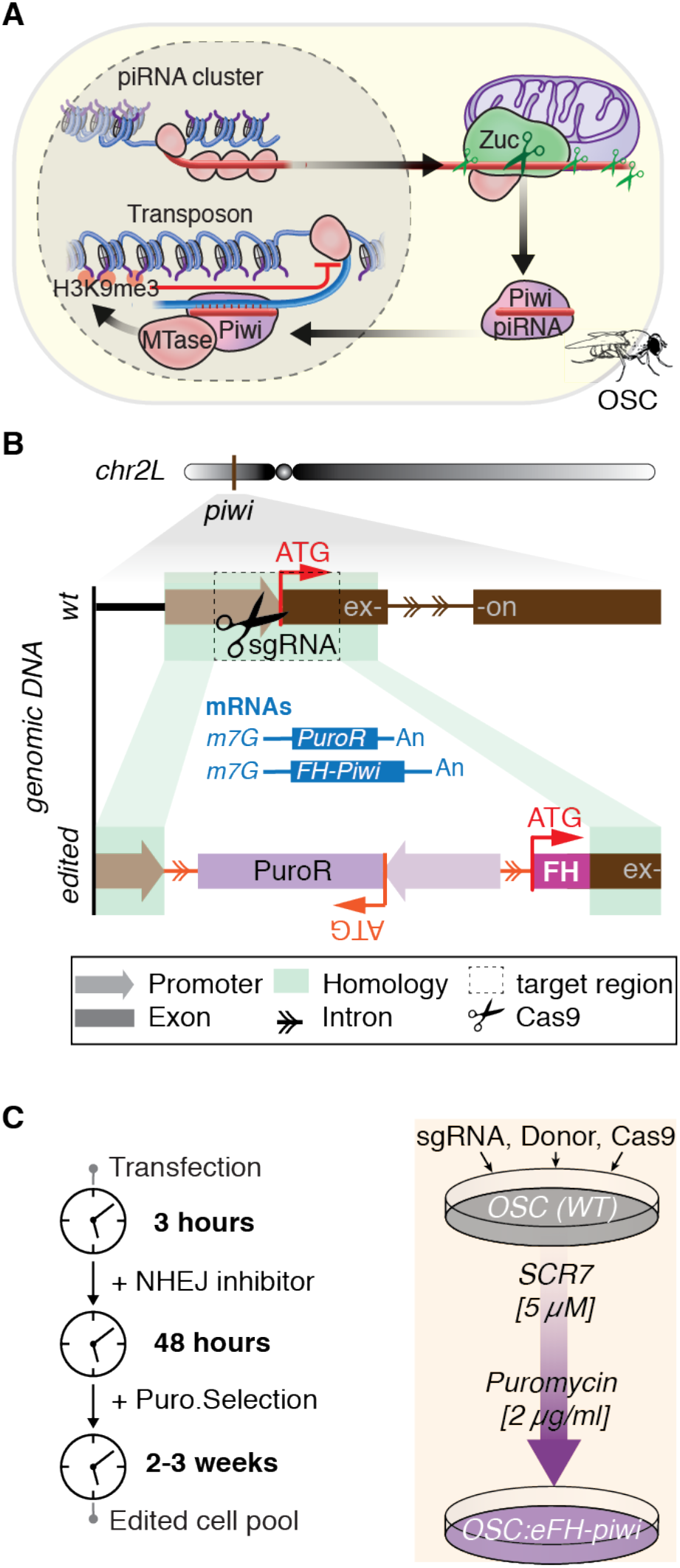
A universal strategy for simple and rapid genomic editing of cell populations. (**A**) *Drosophila* ovarian somatic sheath cells (OSC) represent a unique but delicate model to study Piwi-piRNA mechanism ex vivo. OSC express one of the three *Drosophila* PIWI proteins, Piwi. PiRNAs are generated from long piRNA cluster transcripts by the endonuclease Zucchini (Zuc). Mature Piwi-piRNA silencing complexes transition into the nucleus, recognize nascent transposon transcripts by base-pairing complementarity and induce epigenetic silencing. **(B)** Endogenous tagging of *piwi* in OSC. An sgRNA was designed to target the endogenous *piwi* gene in the vicinity of the start codon (ATG). The donor construct for homologous repair contained a FLAG-HA (FH)-tag and a puromycin resistance gene. The FH-tag was fused in frame with piwi’s open reading frame (ORF) to generate an endogenously N-terminally tagged protein (eFH-). The puromycin resistance was placed into a synthetic intron and transcribed from the opposite genomic strand. The edited allele is designed to express two independent transcripts: The piwi transcript remains under the control of the endogenous promoter and contains an additional intron and a tag. The mature modified mRNA differs from the wt piwi mRNA only by an additional exon-exon junction and the Flag-HA tag. The second transcript is independently generated from the opposite genomic strand and produces an mRNA encoding a puromycin resistance under the control of an Actin promoter. (**C**) Rapid and simple generation of stably edited OSC:*eFH-piwi*. Ovarian somatic sheath cells (OSC) were transfected with an expression plasmid for the sgRNA, the Cas9 endonuclease, and the donor plasmid. Cells were treated with SCR7, an inhibitor of non-homologous end joining (NHEJ) to increase the probability for homologous repair. Antibiotic selection with Puromycin (Puro) was started 48 hours after transfection. After 2-3 weeks, a puromycin resistant cell population has repopulated the dish

### Characterization of the modified cell population

To probe the Puromycin resistant cells for correct genomic insertion of the donor cassette we performed a diagnostic PCR on genomic DNA (gDNA). PCR primers were chosen to detect either the wild type (WT) or the modified allele (eFH-*piwi*) (Fig. 2A). With the intention to generate a universal toolset for endogenous tagging of multiple genes in OSC, we designed a set of primers that recognize a sequence immediately following the splice-donor (SD) or the FH-tag. These universal primers were combined with gene specific primers (ps1_piwi and pas2_piwi) that recognize genomic sequences 5’ and 3’ of the *piwi* homology arms and efficiently detect the wild type (WT) allele. To adapt this strategy to other genes, only the gene-specific primers need to be changed and optimized using WT gDNA. OSC are largely diploid (23), and our genotyping detects a modified and a WT allele in the engineered cells suggesting a heterozygous edit (Fig. 2B).

Next, we tested whether splicing of the introduced intron was accurate and efficient. We placed primers upstream and downstream of the exon-junction to amplify two precise short sequences indicating the spliced transcript in complementary DNA (cDNA). We readily detected the properly spliced transcript in *OSC:eFH-piwi* (Fig. 2C). The un-spliced pre-mRNA was undetectable in cDNA. We would like to note that the context of the splice donor and acceptor sites are crucial for efficient splicing and have been optimized in our donor construct. Thus, we suggest maintaining the nucleotides upstream and downstream of the splice donor and acceptor sites when adapting the construct to target other genomic locations.

**Figure 2.**
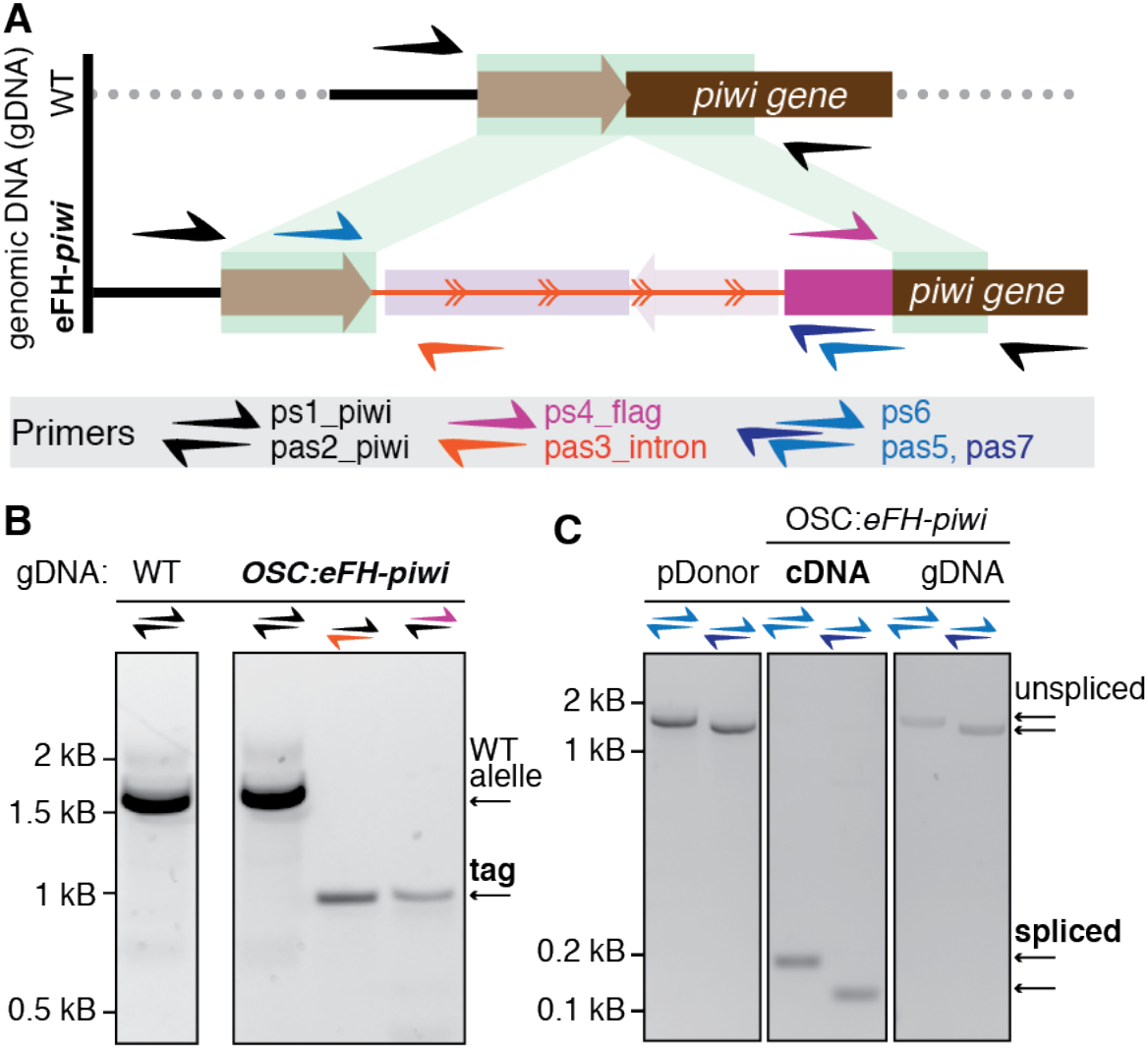
Characterization of the genomic edit and the resulting eFH-piwi transcript. (**A**) Schematic representation of the wild type and the edited piwi allele. The priming sites for universal and gene-specific primers that were used for genotyping and cDNA characterization are indicated. **(B)** Genotyping of OSC:eFH-piwi reveals a heterozygous editing event. PCR with the indicated primers (A) was performed on genomic DNA (gDNA). gDNA from WT OSC was used as control. (C) Characterization of eFH-piwi transcripts indicate accurate splicing of the synthetic intron. PCR was performed on complementary DNA (cDNA) using the indicated primers. Primers were designed to detect the unspliced and spliced transcript (A). The donor plasmid and gDNA served as control for the unspliced transcript.

### Endogenously tagged FH-Piwi (eFH-Piwi) protein maintains wild type expression levels and correct subcellular localization

Next, we tested the expression level of the endogenously tagged Piwi protein (eFH-Piwi). Based on our gDNA analysis that revealed a heterozygous edit, we expected the *OSC:eFH-piwi* to express a tagged and a wild type Piwi protein from the edited and the WT allele respectively. If our engineered allele faithfully maintained the regulation of the endogenous *piwi* and both the mRNA and the protein did not differ in stability, we expected an equal amount of tagged and endogenous Piwi proteins. To quantify the levels of eFH-Piwi and wild type Piwi relative to each other, we separated increasing amounts of cell extract by SDS-PAGE and detected both proteins by western blotting using an endogenous Piwi antibody (Fig. 3A). The FH-tag adds 7.79 kDa to the Piwi protein and allows for discrimination of the tagged and the untagged protein by size. Our quantification revealed that the *OSC:eFH-piwi* expressed both a tagged and a wild type Piwi protein to similar extent.

**Figure 3.**
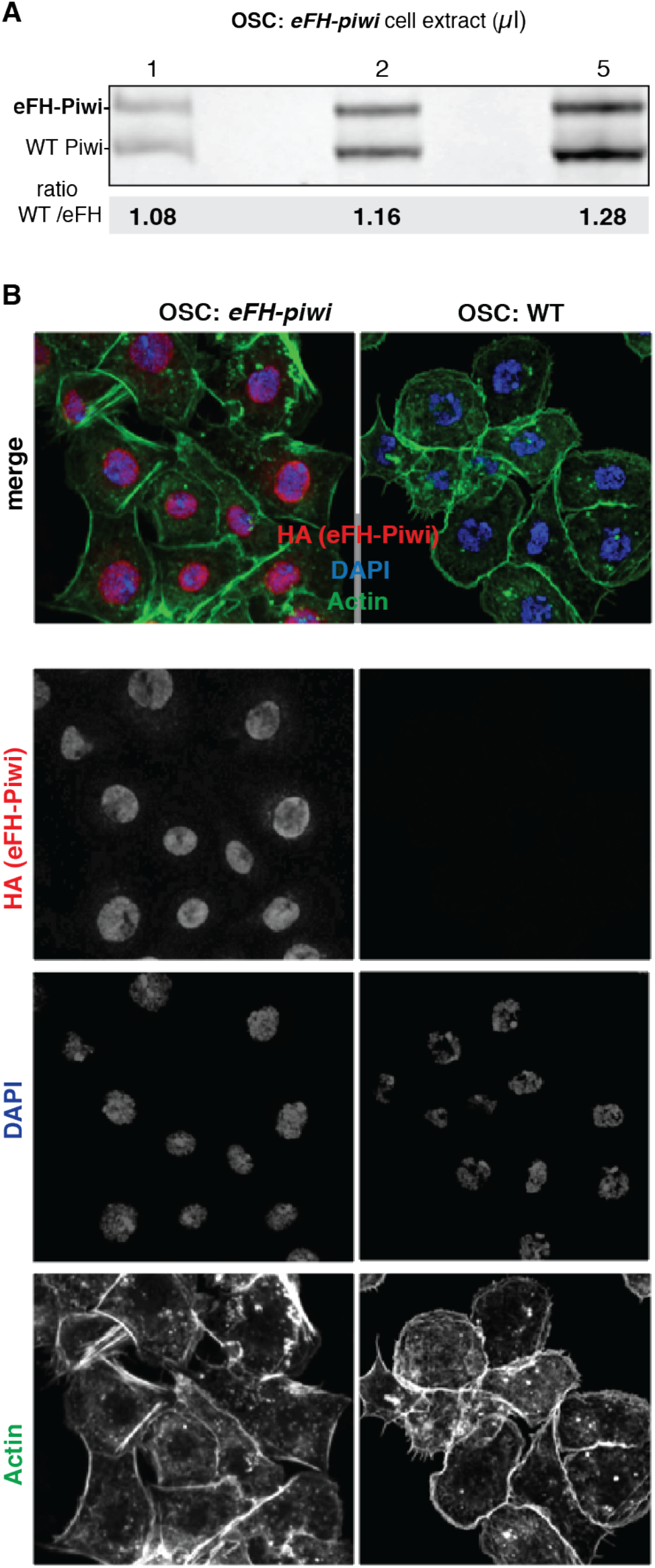
eFH-Piwi protein emulates the expression and subcellular localization of wt Piwi in OSC. (**A**) Heterozygous OSC: *eFH-piwi* expresses WT Piwi and eFH-Piwi protein to similar levels. Wt Piwi and eFH-Piwi were detected with an anti-Piwi antibody. Different amounts of cell extracts were analyzed as indicated. Protein quantification was performed by western blotting using fluorescent antibodies and the LI-COR Odyssey technology for accurate automated quantification. (B) FH-Piwi appropriately localizes to the nucleus of OSC:e*FH-piwi*. The subcellular localization of eFH-Piwi was characterized by immunofluorescence using an anti-HA primary antibody and confocal microscopy. An anti-Tubulin antibody and DAPI were used for cytoplasmic and nuclear counterstain respectively

To evaluate the correct subcellular localization of eFH-Piwi, we performed immunofluorescence analyses using an anti-HA antibody (Fig. 3B). Piwi is known to localize to and function in the nucleus in OSC and in fly ovaries (8, 24, 25). Our results show that the endogenously tagged protein appropriately localizes to the nucleus of *OSC:eFH-piwi* (Fig. 3B).

### PiRNAs associated with eFH-Piwi are comparable to wild type (WT) Piwi-piRNAs by biogenesis signatures, length profile and genomic origin

To directly characterize piRNAs associated with eFH-Piwi and compare them to WT Piwi-piRNAs, we purified eFH-Piwi from *OSC:eFH-piwi* and Piwi from WT OSC, and prepared the associated piRNAs for high-throughput sequencing (Fig. 4). FH-Piwi complexes could be specifically purified from the heterozygous OSC:*eFH-piwi* using a specific anti-FLAG antibody under stringent wash conditions (Fig. 4A). Associated piRNAs were extracted and cDNA libraries were generated for Illumina sequencing (19). For accurate quantification of small RNA reads, we included ten unique molecular identifiers (UMI) (26) in the ligated adapters before cDNA preparation and PCR amplification (Fig. 4B). These UMIs enable elimination of PCR-duplicates during data analyzes and thus ensure an unbiased representation of the sampled small RNA population.

Piwi-piRNAs are produced by the phased action of the Zucchini-processor complex that generates a characteristic preference for Uridine (U) in the first position of the mature piRNAs (9, 10, 18). Additional preferences for Uridine can be observed one piRNA length upstream and downstream (position −26 and +26) indicating proceeding and preceding piRNAs in a metagene analysis (18). These processing signatures can be readily observed for WT Piwi-piRNAs in OSC and for piRNAs associated with eFH-Piwi (Fig. 4C). Uniquely mapping piRNA associated with WT Piwi or with eFH-Piwi were aligned at their first position and the genomic interval was extended to include one piRNA length upstream and one piRNA-length downstream of the observed molecules. The relative frequency of all four nucleotides was calculated for each position across a 100 nucleotide (nt) window and revealed the characteristic phased 1U-signature. These results suggest that eFH-Piwi, like WT Piwi, is fueled with piRNAs that are generated by the ZUC-processor complex.

Next, we compared the length distribution of piRNAs associated with WT Piwi or with eFH-Piwi. The length profiles of piRNAs are characteristic for their associated PIWI protein and have been suggested to reflect a footprint of the PIWI protein during 3’ end formation (24, 27, 28). Like WT Piwi-piRNAs, eFH-Piwi-piRNAs exhibit a preferred length of 25-26nt (Fig. 4D).

Finally, we tested whether both piRNA populations originate from the same genomic regions. Piwi-piRNAs originate to a large part from precursors with antisense complementarity to transposons and other repetitive elements that can be annotated by Repeatmasker (rmsk) (24). Results from our analysis show that more than half of the piRNAs associated with WT and eFH-Piwi contain sequences that are antisense to genomic repeats (Fig. 4E).

**Figure 4.**
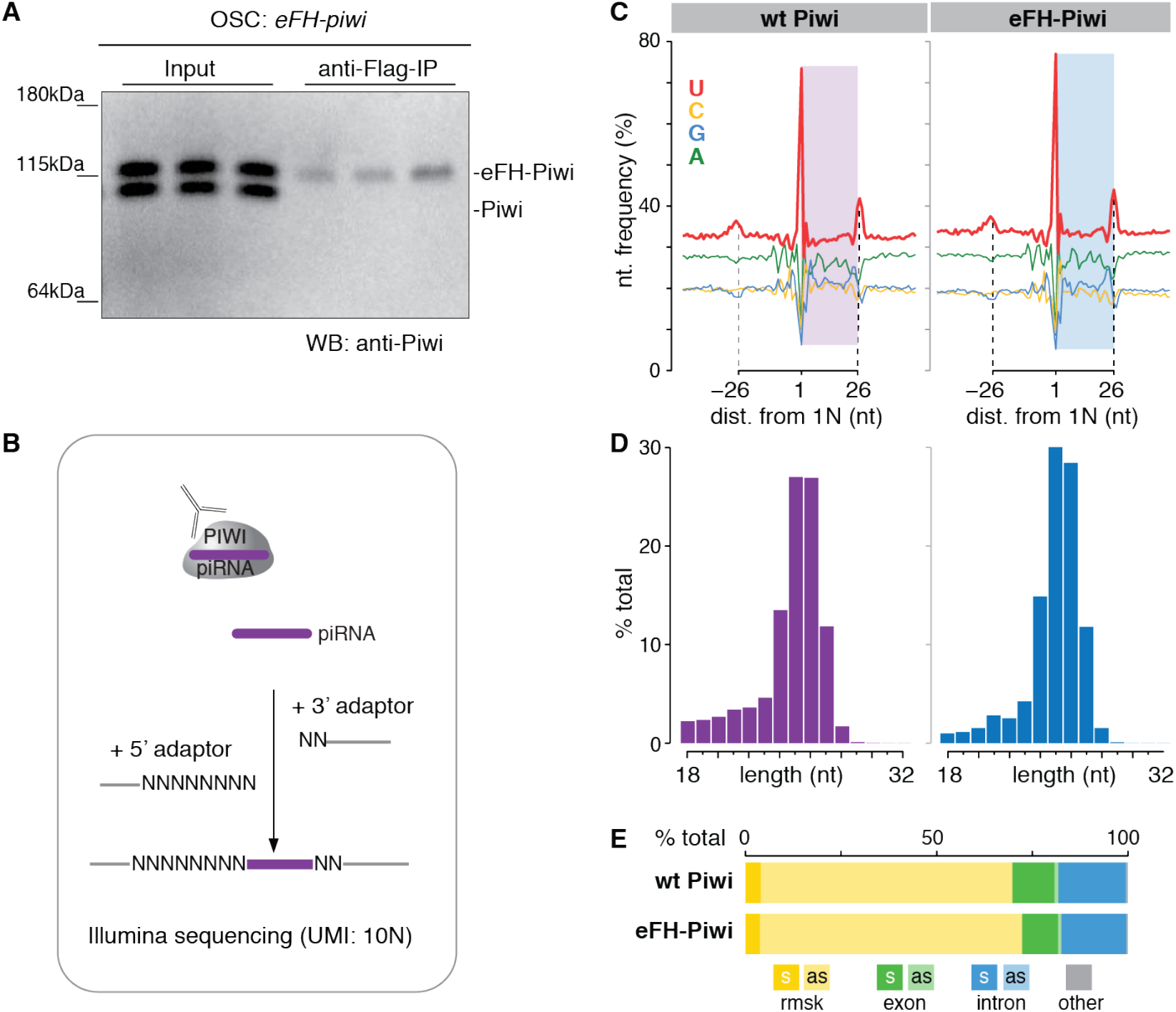
eFH-Piwi associates with piRNAs to form mature piRNA silencing complex. **(A)** eFH-Piwi was specifically immunopurified (IP) from OSC: *eFH-piwi* using an anti-Flag antibody. **(B)** Small RNAs were extracted from the purified Piwi-piRNA complexes and prepared for Illumina sequencing. 3’ and 5’ adaptors were sequentially ligated to the small RNAs before reverse transcription and PSC amplificaion. A total of 10 unique molecular identifiers (UMI: 10N) was accommodated in the ligated adaptors and allowed for removal of PSC duplicates during analyses. (**C**) PiRNAs associated with eFH-Piwi exhibit the same phased III-signatures as WT Piwi-piRNAs, indicating biogenesis by the Zuc-processor complex. Metagene analysis of uniquely mapping piRNAs aligned at their 5’ end across an extended genomic interval. The observed piRNA population is indicated as colored box. Nucleotide frequencies are shown across a 100 nt interval. Both piRNA populations show the characteristic patterns of phased processing by the Zucchini processor complex indicated by a preference for Uridine in the first position (1U), and both one piRNA length upstream (−26) and one piRNA length downstream (26) of the observed piRNAs. (**B**) Length profiles of piRNAs associated with eFH-Piwi and WT Piwi in nucleotides (nt). Both piRNA populations show a length distribution characteristic for Piwi-piRNAs. (**C**) eFH-Piwi-piRNAs, like WT Piwi-piRNAs are enriched for sequences that are antisense to annotated repeats (rmsk). More than half of either piRNA population represents sequences with antisense complementarity to transposons and other repeat elements (repeat masker, rmsk). The orientation with respect to the matching feature is indicated (sense, s; antisense, as)

Taken together, results from our molecular and computational analyses show that eFH-Piwi emulates the expression level and subcellular localization of WT Piwi and forms complexes with piRNAs that are indistinguishable from wild type piRNAs. Our approach combines precise genome editing with simple antibiotic selection to generate stably edited cells that express an endogenously tagged protein within a few weeks. Our strategy bypasses the need for selection of edited cell clones, which is laborious and does not work effectively for all cell types. Overall, we aim to share the strategy and reagents that enable the rapid establishment of endogenously tagged proteins and provide means to add high-affinity and high-specificity tags for biochemical and biological experiments.

## DISCUSSION

While in the case of Piwi, specific and sensitive antibodies for detection and purification are available (24), such tools are often missing especially for germline specific proteins. Generation of effective antibodies is laborious, costly, and often unsuccessful, even with a sincere effort. The addition of a Flag-HA (FH) tandem tag to *piwi* enabled us to increase stringent washes during Piwi-piRNA purification and generated a relevant biological tool to study the Piwi-only pathway in OSC.

Conventional strategies of endogenous tagging often involve selection, characterization, and growth of individual cell clones, a laborious and time-consuming procedure. Furthermore, clonal selection establishes a novel cell line with unique characteristics and requires the analyses of multiple clonal lines and rescue experiments to confidently exclude changes due to clonal selection. Our approach capitalizes on the advantages of endogenous tagging while eliminating the need for clonal selection by the alternative of antibiotic selection.

Accommodating the antibiotic resistance in an optimized synthetic intron provides an opportunity to not only edit the beginning or end of a gene, but also to modify gene-internal sequences. Our donor cassette could be modified to generate mutations or deletions within a gene body. Such an approach could be used to obliterate or mimic sites of post-translational modifications, change enzymatic activities, or engineer non-coding RNAs. Overall, our method describes a simple and rapid technique to generate edited cell pools with facility cell biological experiments.

